# A pipeline for rapidly generating genetically engineered mouse models of pancreatic cancer using in *vivo* CRISPRCas9 mediated somatic recombination

**DOI:** 10.1101/398347

**Authors:** Noboru Ideno, Hiroshi Yamaguchi, Takashi Okumara, Jonathon Huang, Mitchel J. Brun, Michelle L. Ho, Junghae Suh, Sonal Gupta, Anirban Maitra, Bidyut Ghosh

## Abstract

Genetically engineered mouse models (GEMMs) that recapitulate the major genetic drivers in pancreatic ductal adenocarcinoma (PDAC) have provided unprecedented insights into the pathogenesis of this lethal neoplasm. Nonetheless, generating an autochthonous model is an expensive, time consuming and labor intensive process, particularly when tissue specific expression or deletion of compound alleles are involved. In addition, many of the current PDAC GEMMs cause embryonic, pancreas-wide activation or loss of driver alleles, neither of which reflects the cognate human disease scenario. The advent of CRISPR/Cas9 based gene editing can potentially circumvent many of the aforementioned shortcomings of conventional breeding schema, but ensuring the efficiency of gene editing *in vivo* remains a challenge. Here we have developed a pipeline for generating PDAC GEMMs of complex genotypes with high efficiency using a single “workhorse” mouse strain expressing Cas9 in the adult pancreas under a *p48* promoter. Using adeno-associated virus (AAV) mediated delivery of multiplexed guide RNAs (sgRNAs) to the adult murine pancreas of *p48-Cre; LSL-Cas9* mice, we confirm our ability to express an oncogenic *Kras* ^*G*^*12*^*D*^ allele through homology-directed repair (HDR), in conjunction with CRISPR-induced disruption of cooperating alleles (*Trp53, Lkb1* and *Arid1A*). The resulting GEMMs demonstrate a spectrum of precursor lesions (pancreatic intraepithelial neoplasia [PanIN] or Intraductal papillary mucinous neoplasm [IPMN] with eventual progression to PDAC. Next generation sequencing of the resulting murine PDAC confirms HDR of oncogenic *Kras*^*G*^*12*^*D*^ allele at the endogenous locus, and insertion deletion (“indel”) and frameshift mutations of targeted tumor suppressor alleles. By using a single “workhorse” mouse strain and optimal AAV serotype for *in vivo* gene editing with combination of driver alleles, we have created a facile autochthonous platform for interrogation of the PDAC genome.

## INTRODUCTION

Pancreatic ductal adenocarcinoma (PDAC) is one of the most lethal cancers known, due to the absence of reliable early diagnosis and the absence of effective therapeutic regimens [1]. It is currently the third leading cause of cancer worldwide and is projected to become second leading cause of cancer in the United States by 2030. The median 5-year survival rate of PDAC is only 9% [2], and underscores the need for developing new approaches in understanding the molecular mechanisms underlying the disease.

The advent of next generation sequencing (NGS) and its application as part of large publicly funded efforts such as The Cancer Genome Atlas (TCGA) and International Cancer Genome Consortium (ICGC) has identified a huge catalogue of genes that are altered in PDAC [3, 4], although in many instances, particularly with low frequency mutations, the functional consequences of the alterations remain unknown. The definitive evidence for gene perturbation in carcinogenesis is typically obtained via autochthonous models, which in the context of PDAC, is exemplified by expression of a mutant *Kras* allele in the pancreas alongside one or more cooperating mutations [5]. First developed in 2003, PDAC GEMMs have dramatically accelerated our understanding of the molecular pathogenesis of this neoplasm, including how the genetic background of the cancer impacts the tumor microenvironment (TME) and resulting immune response [6-8]. In contrast to cell line or patient-derived xenograft (PDX) models, autochthonous models harbor the precursor lesions observed in the multistep progression of PDAC, as well as the stroma-dense TME observed in the corresponding human disease [9-11].

Although autochthonous models have been instrumental in our understanding the molecular underpinnings of PDAC, the conventional *Cre*-based recombinatorial models do not faithfully mimic adult onset human PDAC due to their inherent limitation of inducing *Ras* activation and cooperating gene alteration(s) in every epithelial cell of the pancreas during development. In some respects, the conventional models are more analogous to “familial” PDAC where every cell in the pancreas is predisposed to neoplasia due to a germline anomaly (for example, *CDKN2A/p16*), and cancers arise due to stochastic bi-allelic gene inactivation [12, 13]. In addition to this conceptual limitation, conventional *Cre*-mediated recombinatorial models are also hindered by time, labor and expenses required for generating appropriate numbers of colonies, especially with complex genotypes where the “pups” of interest might only be a minor fraction of the litter. A timeframe of 12-18 months for completing the breeding of complex mouse genotypes is not unusual in the field. Novel platforms that bypass the pancreas-wide embryonic expression of mutant *Kras* and loss of cooperating alleles, while also greatly improving the efficiency of model generation would be a significant advancement in the preclinical arena.

Recently, the RNA-guided endonuclease *Cas*9 from microbial type II CRISPR (clustered regularly interspaced short palindromic repeat) system has emerged as a powerful tool for genome engineering in mammalian cells [14-16], including somatic cell editing in a variety of mouse organs that includes brain, lung, liver and pancreas [17-21]. CRISPR/Cas9 can be targeted to the desired genomic loci by using programmable 20-bp single guide RNAs (sgRNAs) to generate DNA double stranded breaks which induce genome editing via one of the two DNA damage repair pathways: non homologous end-joining (NHEJ) resulting in insertion-deletion (INDELs), or homology directed repair (HDR) resulting in precise sequence substitution in the presence of a repair template [22-25]. The multiplexing abilities of gene editing by combining multiple sgRNAs to the Cas9 provides this programmable nuclease system with a unique advantage for conducting *in vivo* combinatorial gene editing, especially in the context of developing autochthonous cancer models [21]. There have been several approaches used for delivering CRISPR/*Cas*9 *in vivo* in murine pancreatic cells, including lentivirus and plasmid based transfection [26, 27], although many of these vectors are hindered by low efficiency of multiplexed gene editing and risk of on-target recombination in unintended cell types or tissues [28]. In recent years, adeno-associated virus (AAV) vectors, in particular, have been increasingly utilized for *in vivo* delivery approaches of sgRNAs due to their efficiency, lack of inherent pathogenicity and strong safety profile [29-32].

In order to generate a “workhorse” model of genetically engineered mice that develop PDAC with a variety of complex genotypes, we have developed a simple injection method by employing AAV-based delivery of sgRNAs cargo to the pancreas of adult mice, where *Cas*9 activity is restricted to the p48-expressing acinar compartment [33]. Our model overcomes several challenges of the current *Cre-*based models, including the introduction of mutant alleles in a subset of adult pancreatic epithelial cells (rather than pancreas-wide embryonic expression), and the use of a single “workhorse” mouse genotype (*p48*-Cre; LSL-*Cas*9) from which cancers harboring multiple complex genotypes can be readily generated through somatic gene editing, thus greatly increasing efficiency. Importantly, in contrast to some of the recently described CRISPR/*Cas*9-mediated PDAC models that require an oncogenic *Kras*^G12D^ allele in the pancreas on which cooperating mutations are then introduced by CRISPR/*Cas*9 [26, 27], our method dispenses with the need for a constitutive *Kras*^G12D^ allele, by including a mutant *Kras*^G12D^ template in the delivery cargo that recombines with, and replaces, the endogenous wild type allele through HDR.

## MATERIALS AND METHODS

### Generation of pancreas specific *Cre*-dependent Cas9 mice

Conditional *Cas9* mice (*Rosa26-LSL-Cas9*) was obtained from the Jackson Laboratory and was crossed with pancreas specific *p48-Cre* mice in order to generate experimental *p48-Cre; LSL-Cas9* mice. The following primers were used to genotype *Cas9* (Cas9_Common-S 5’-AAGGGAGCTGCAGTGGAGTA-3’; Cas9_Wild-AS 5’-CCGAAAATCTGTGGGAAGTC-3’; Cas9_Mut-AS 5’-CGGGCCATTTACCGTAAGTTAT-3’) and *p48-Cre* (ptf1a_Fw 5’-AACCAGGCCCAGAAGGTTAT-3’; ptf1a_Rv 5’-TCAAAGGGTGGTTCGTTCTC-3’; ptf1a_Cre_Fw 5’-ATAGGCTACCTGGCCATGCCC-3’;ptf1a_Cre_Rv5’-CGGGCTGCAGGAATTCGTCG-3’).

### CRISPR sgRNA design

CRISPR sgRNAs for *Kras*, *Trp53* and *Lkb1* have been previously described [19]. Genomic sequence for *Arid1a* gene was downloaded from UCSC genome browser and sgRNA cassette was generated using the CRISPR design tool (http://crispr.mit.edu).

### AAV vector plasmids

The AAV-KPL plasmid (AAV: ITR-U6-sgRNA (*Kras*)-U6-sgRNA (*p53*)-U6-sgRNA (*LKB1*)- pEFS-RLUC-2A-Cre-shortPA-KrasG12D_HDRDonor-ITR) was obtained from Addgene. AAV-KPLδCre was generated by removing the Cre sequence from AAV-KPL by digesting with Nhe1 and HindIII followed by ligation. AAV-KPδCre was generated by digesting AAV-KPLδCre with Kpn1 and Xba1 and followed by ligation. AAV-KδCre was made by digesting AAV-KPLδCre with BamH1 and Kpn1, and followed by ligation. To construct AAV-KPAδCre, first the sgRNA for mouse *Arid1a* exon3 was cloned into the pX459 (pSpCas9 (BB)-2A-Puro) vector, then the whole segment containing the U6 promoter, *Arid1a* sgRNA with primers containing Xba1 and Kpn1 sites was PCR amplified. Finally, AAV-KPLδCre was digested with Xba1 and Kpn1 to release the *Lkb1* CRISPR containing unit following which the entire amplicon containing *Arid1a* CRISPR was ligated to generate AAV-KPAδCre. AAV-KAδCre was made by digesting AAV-KPAδCre with BamH1 and Xba1 and re-ligated. All of the AAV vector constructs were sequence verified before virus production. All vector constructs are readily available from the authors upon request (the schematic of the constructs are shown in **Supplementary Fig. 1A**).

### AAV vector production

Human Embryonic Kidney 293T (HEK293T) cells were used for AAV production. HEK293T cells were maintained in Dulbecco’s Modified Eagle Medium (DMEM, Life Technologies) supplemented with 10% fetal bovine serum (FBS, Atlanta-Biologicals) and 1% penicillin and streptomycin (Life Technologies). Cells were grown as adherent cultures in 5% CO2 at 37 °C on 15 cm cell culture plates.

Viruses were generated by linear polyethyleneimine (PEI)-mediated triple transfection. The PEI was added to a DNA mixture of pAAV2/8, pAAV2/5, or pAAV2/6 *rep/cap* plasmid (10 µg/plate), transgene plasmid (10 µg/plate), and pXX6-80 helper plasmid (20 µg/plate), incubated for 30 min at room temperature and added to poly-L-lysine-coated plates of 90% confluent HEK293T cells. The cells were harvested 48 h post-transfection and lysed by three cycles of freeze/thaw. The cell lysate was treated with 4 U/ml Benzonase (Sigma-Aldrich) for 40 min at 37°C, and then separated through an iodixanol gradient (15/25/40/54%) in Beckman Ultra-Clear Quick Seal Tubes (Beckman Coulter, Brea, CA). After centrifugation at 48,000 rpm for 1.75 h in a 70Ti rotor, the virus was extracted from the 40% layer. Viruses were further purified through anion exchange chromatography (Q Sepharose, Pall) followed by concentration and buffer exchange through Amicon Ultra 4 (Millipore) into GB-PF68 (50 mM Tris, pH 7.6, 150 mM NaCl, 10 mM MgCl_2_, 0.001% Pluronic F68). Virus titers were determined by qPCR using SYBR green (Life Technologies) and primers against the Rluc portion of the transgene cassette (forward: GCCTCGTGAAATCCCGTTAGTA, reverse: GCATTGGAAAAGAATCCTGGGTCC) on a BIO-RAD CFX96.

### Surgical procedure and direct injection

100 μl of PBS containing AAV-CRISPR sequences was injected into the pancreatic tail using 29 G needle (BD Biosciences) through left subcostal laparotomy under inhaled anesthesia. Leakage was not observed in any mice during the procedure. After the injection, the pancreas and spleen was carefully returned into the peritoneal cavity and the abdomen was closed using 4-0 Vicryl (Ethicon) and skin staplers. All the mice were monitored at least once a week by abdominal palpation and abdominal ultrasound to investigate the entire pancreas was performed every 4 weeks using the Vevo 2100 (Fujifilm). Moribund mice with pancreatic tumors were euthanized by cervical dislocation and carbon dioxide inhalation according to the IACUC protocol. The assessment of primary tumor, intraperitoneal dissemination and distant metastases performed were based on both gross findings at necropsy and by histopathological examination. Any surviving mice at 100 days (∼ 3 months post injection) were then euthanized and the Pancreata harvested for histopathological examination.

### T7 Endonuclease Gene editing assay

Genomic DNA was isolated from tumor samples using the Qiagen genomic DNA isolation kit. Subsequently ∼400bp of genomic fragment encompassing the CRISPR targeted cleavage site was PCR amplified with high fidelity Taq polymerase. Approximately 200ng of purified PCR product was used for each T7 endonuclease assay. Briefly, 2ul NEBuffer 2, 200ng of purified PCR product and dH2O was added to a total of 19ul in a PCR tube. The hybridization reaction was run as follows: 5min, 95C; ramp down to 85C at -2C/s; ramp down to 25C at -0.1C/s; hold at 4C. We then added 1ul (10U) T7 endo I and incubated at 37C for 15min. The reaction was stopped by adding 2ul of 0.25M EDTA and loaded immediately on a 1.5% agarose gel.

### Immunofluorescence

Pancreata were harvested from the moribund mice due to tumor burden at different time point with maximum harvesting time around ∼100 days. Pancreata were fixed in 4% paraformaldehyde, followed by standard paraffin embedding. Tissue was then cut into 5uM sections. Incubations with primary antibodies were performed overnight at 4^0^C using standard techniques in PBS containing 0.2% Triton and 10% FBS using the following antibodies or probes: mouse anti-E-Cadherin (BD Transduction 610181, 1:50); rabbit anti-Sox9 (Millipore AB5535, 1:1000); hamster anti-Muc1 (Thermo Fisher HM-1630-F, 1:100); mouse anti-Arid1a (Santa Cruz Biotechnology sc-32761, 1:100); mouse anti-p53 (Santa Cruz Biotechnology sc126, 1:50) and rabbit anti-LKB1 (Abcam ab185734 1:100). For immunofluorescence, secondary antibodies were obtained from Abcam and used at 1:300 dilution; samples were mounted with Fluorescence mounting media (Dako, S3023) and imaged on the Olympus Confocal Microscope.

### Whole Exome Sequencing

Whole exome sequencing of harvested murine tumors was performed at Johns Hopkins University Next Generation Sequencing (NGS) Core Facility. Briefly, the Agilent SureSelect Mouse exome sequencing kit was used to prepare sequencing libraries and capture annotated exonic sequences. The resulting libraries were sequenced on a HiSeq 2500, and after quality control checks the fastq files were aligned to genome build mm10 with bwa mem 0.7.7. Picard tools 1.118 were used to remove duplicate reads and GATK 3.6 was used for base recalibration. To call somatic variants, a reference sequence panel of normals was first constructed from C57BL mouse strains, and Mutect2 was used with this panel. Annotation was done with snpEFF and snpSIFT.

## RESULTS

### Strategy for virally delivered CRISPR-mediated genome editing in the adult pancreatic epithelium

As a prelude for *in vivo* gene editing studies we first performed direct injection of AAV containing reporter alleles into the mouse pancreas. To test the differential efficacy of distinct AAV serotypes infecting pancreatic epithelial cell, we injected AAV6-Cre-GFP and AAV8-Cre-GFP into the adult mouse Pancreata (**Supplementary Figure.1B, C**) which confirmed the superiority of AAV8 in infecting epithelial cells as evidenced by extent of GFP expression. Henceforth, all of our AAV vectors were made using AAV8. Further, in order to avoid spurious *Cre*-dependent recombination in the non-epithelial pancreatic tissue including the peritoneum from the injected virus, we deleted the *Cre* sequence from the original AAV construct, AAV-KPL (**Supplementary Figure.1A**). Instead, we generated *p48-Cre*^*+/-*^; *LSL-Cas9* mice as the “workhorse” mouse where Cre recombinase is only expressed in the adult pancreatic epithelium, specifically acinar cells.

### Multiplexed CRISPR/Cas9 mediated somatic genome-editing results in autochthonous PDAC harboring diverse genotypes

A set of six independent AAV8 vectors were generated for injection into the pancreas of *p48-Cre*^*+/-*^*;LSL-Cas9* mice, containing the following sgRNAs: (i) *AAV-LacZ* (control) (ii)*AAV-Kras* (which included the *Kras* ^*G*^*12*^*D*^ HDR template and henceforth designated as “K”, (iii) *AAV-Kras*, *Trp53* (henceforth designated as “KP”), (iv)*AAV-Kras; Arid1a* (henceforth designated as “KA”), (v) *AAV-Kras; Trp53; Lkb1* (henceforth designated as “KPL”) and (vi) *AAV-Kras;Trp53;Arid1a* (henceforth designated as “KPA”). The schematic of the model development and post-harvest correlative assays are illustrated in **Figure 1A**. The pancreatic development was monitored in real time by ultrasound as shown in **Figure 1B**. The survival of the mice injected with the six vectors is depicted in **Figure 1C**. Excluding the control “LacZ” mice, the mice with “K” and “KA” had the most indolent natural history with no mortality observed at ∼100days (at which point these mice were euthanized). In contrast, “KP”, “KPA” and “KPL” injected mice had comparable median survival with progression to invasive neoplasia. In the “KPL” injected nice, ultrasound examination demonstrated readily visible tumors as early as eight weeks after injection and these were apparent 1-2 weeks later in “KP” and “KPA” injected mice.

**Figure 1.**
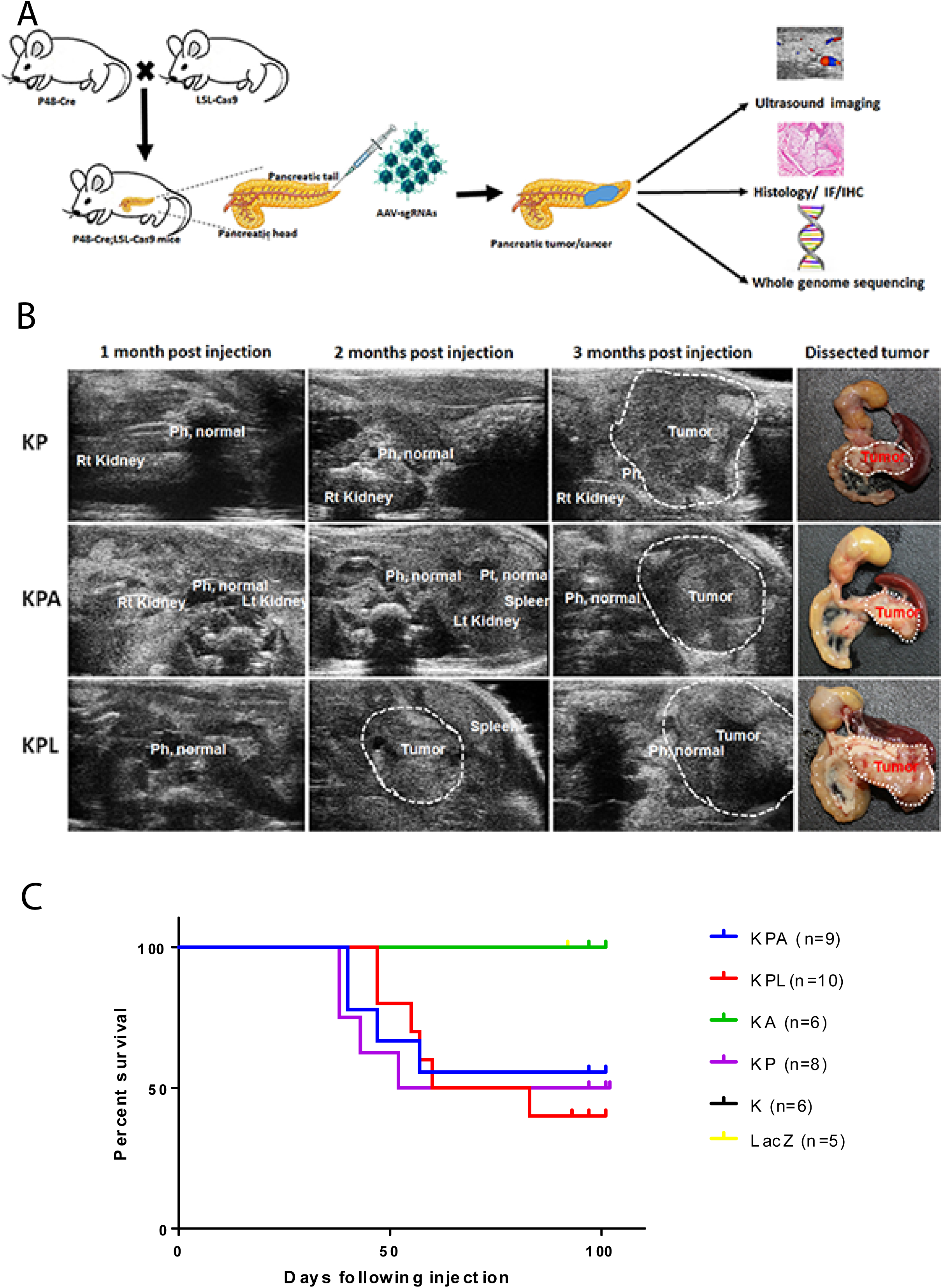
Generation and natural history of CRISR/Cas9 induced genetically engineered mouse models of pancreatic cancer. **A**. Schematic of experimental procedure showing the direct injection of AAVs containing sgRNAs into the tail of *p48*-Cre; LSL-*Cas9* mice. Serial ultrasound imaging was performed to follow tumor induction for ∼3 months following injection, at which time point the mice were euthanized and the pancreata were harvested for histopathological evaluation and correlative studies, **B**.Non-invasive monitoring of pancreatic tumor formation by ultrasound imaging at regular interval following viral injection. Note the normal pancreas head (Ph) at 3 months post injection while the pancreatic tail harbors a large tumor. Relative position of the pancreas with respect to kidney and spleen are depicted on the ultrasound image. **C**. The natural history of various tumor genotypes generated in this study – LacZ (control), “K”, “KA”, “KP”, “KPA”, and “KPL” is show via Kaplan-Meier survival curve. Any remaining mice alive at 100 days post-injection were euthanized and underwent necropsy. Please see text for details of each genotype.

Representative histopathology photomicrographs from the mice injected with various AAV constructs are shown in **Figure 2**. In mice that succumbed to invasive carcinomas, we have captured examples of murine pancreatic intraepithelial neoplasia (mPanIN) in the surrounding pancreas, to demonstrate that the carcinomas arose on a backdrop of what is widely accepted as the most common precursor lesion in human PDAC. In “K” and “KA” injected mice, we only observed focal mPanIN lesions of various grades, in the absence of invasive carcinomas (**Figure 2 A-D**), which is also reflected in their survival. Notabley, surrounding histologically normal pancreatic parenchyma was readily seen in these mice even at ∼ 100 days, distinct from the pancreas-wide activation of *Cre* in conventional models where there is progressive replacement of acinar tissue by precursor lesions throughout the organ. In contrast to mice injected with “K” and “KA” viruses, the remaining cohorts developed invasive cancers, as seen with “KP” (**Figure 2 F**), “KPA” (**Figure 2 H**) and “KPL” (**Figure 2 J**) respectively. Of note, we also readily observed examples of acinar ductal metaplasia (ADM) in the background parenchyma and typically adjacent to mPanINs, as these are considered the “precursor to the precursor” in the multistep model of murine PDAC pathogenesis (**Figure 2 E**, **G** and **I**). Further, the “KPL” injected mice were distinguished by the presence of cystic precursor lesions (Intraductal papillary mucinous neoplasms or IPMNs, illustrated in **Supplementary Figure 2**), a feature that has been previously reported in conventional *Cre* models of *Lkb1* deletion. The resulting IPMNs expressed the apomucin MUC1 on their surface, consistent with a so-called pancreato-biliary subtype of differentiation (**Supplementary Figure 3**)

**Figure 2.**
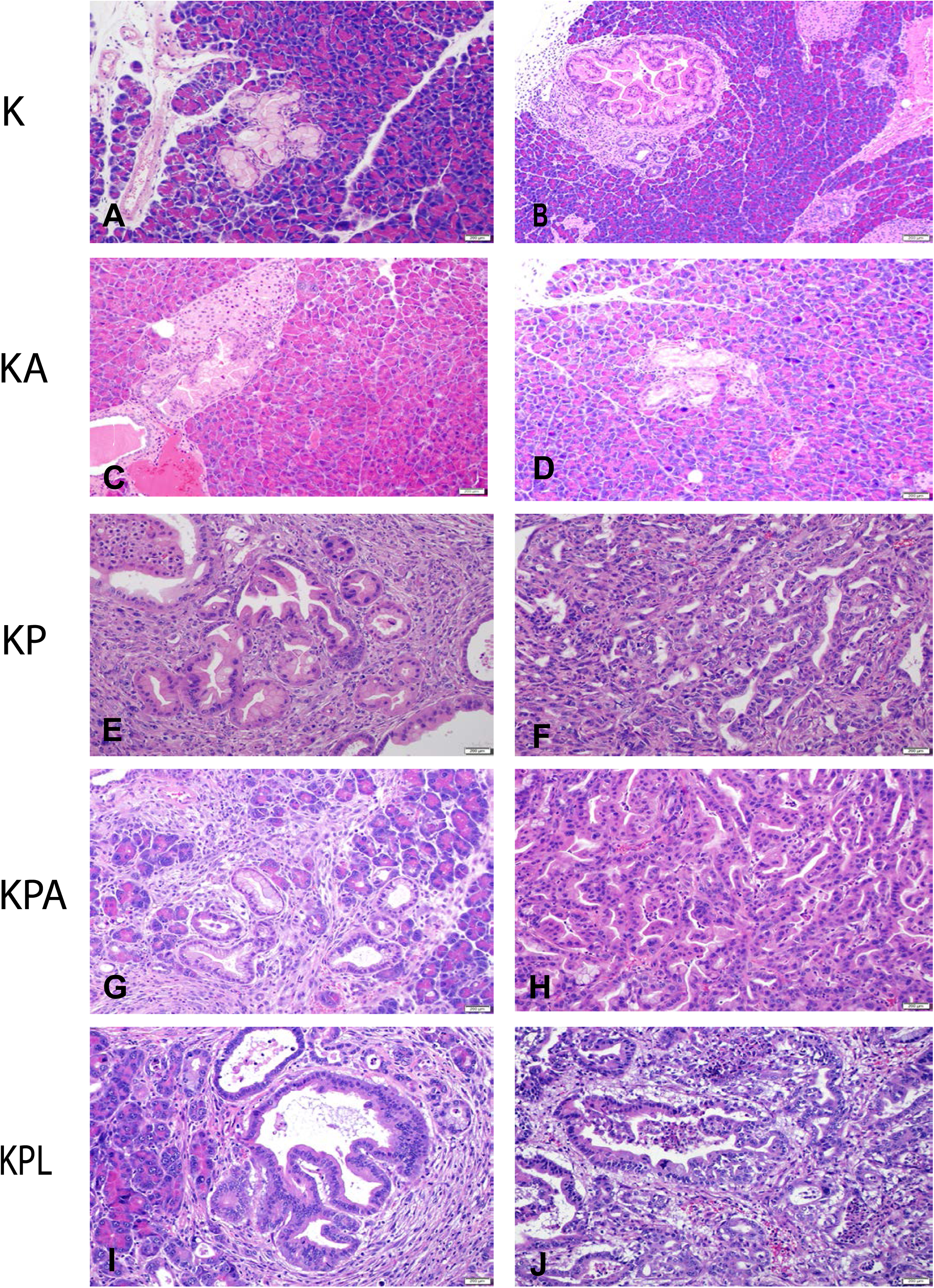
Histopathology of CRISPR-Cas9-induced pancreatic precursor lesions and invasive cancers in *p48*-Cre; LSL-*Cas9* mice. **A**.Low-grade murine pancreatic intraepithelial neoplasia (mPanIN) lesion on an otherwise uninvolved pancreatic parenchyma in an AAV-*Kras* (“K”) mouse. No invasive cancers were seen at ∼100 days. **B.**High-grade murine mPanIN lesion on an otherwise uninvolved pancreatic parenchyma in an AAV-*Kras* (“K”) mouse. No invasive cancers were seen at ∼100 days. **C-D.** Low-grade murine mPanIN lesion on an otherwise uninvolved pancreatic parenchyma in AAV-*Kras; Arid1a* (“KA”) mice. No high-grade mPanIN or invasive cancers were seen at ∼100 days. **E**. Entrapped high-grade murine mPanIN lesions with poorly differentiated (sarcomatoid) PDAC in the background in an AAV-*Kras, Trp53* (“KP”) mouse. **F**. Invasive adenocarcinoma (PDAC) arising in an AAV-*Kras, Trp53* (“KP”) mouse. **G**.Low-grade murine mPanIN lesions and acinar ductal metaplasia (ADM) in a pancreas with changes of chronic pancreatitis in an AAV-*Kras, Trp53, Arid1A* (“KPA”) mouse. **H.**Invasive adenocarcinoma (PDAC) arising in an AAV-*Kras, Trp53, Arid1A* (“KPA”) mouse. **I.**High-grade murine mPanIN lesions and acinar ductal metaplasia (ADM) in a pancreas with changes of chronic pancreatitis in an AAV-*Kras, Trp53, Lkb1* (“KPL”) mouse. **J.**Invasive adenocarcinoma (PDAC) arising in an AAV-*Kras, Trp53, Lkb1* (“KPL”) mouse.

Histopathological examination of the invasive carcinomas at low magnification often showed normal pancreatic parenchyma adjacent to a region of invasive carcinoma mirroring the “segmental” nature of disease in the human pancreas (and distinct from the multifocal nature of disease seen in the conventional *Cre* models) (**Supplementary Figure 4A**). While adenocarcinomas were seen with all three cancer forming genotypes (“KP”, “KPA” and “KPL”) (**Figure 2**), the proportion of adenocarcinomas versus poorly differentiated (sarcomatoid) tumors varied (**Supplementary Figure 4B**), with “KPL” and “KPA” resulting predominantly in adenocarcinomas and “KP” in poorly differentiated lesion. These variations in the proportion of histological subtypes for individual genotypes have been well documented with *Cre*- based models [34], so was not unexpected. In addition to the primary tumor, metastatic adenocarcinomas were observed in a subset of “KPL” and “KPA” injected mice, specifically in the liver and lung (**Supplementary Figure 5**).

### Correlative studies in CRISPR/Cas9 generated PDAC models

A series of correlative protein expression and sequencing studies were conducted on the harvested tissues from the CRISPR/Cas9 generated PDAC lesions. For example, we confirmed the loss of expression of p53, Arid1a and Lkb1 within the neoplastic cells of tumors harvested from “KPL” and “KA” and virus injected pancreata, respectively (**Figure 3**). Although “KA” injected mice did not form invasive carcinomas, partial to complete loss of expression of Arid1a protein was demonstrable in the ADM and mPanIN lesions that resulted in the pancreata (identified based on Sox9 expression) consistent with successful somatic editing of *Arid1*a loci (**Supplementary Figure 6**). As previously noted “KP” mice developed multiple examples of poorly differentiated (sarcomatoid) carcinomas, and we confirmed the epithelial origin of these cancers by the expression of cytokeratin 19 in the primary tumors (data not shown). Of note, these poorly differentiated PDAC were characterized by loss of membrane E-cadherin and co expression of the mesenchymal marker vimentin, suggesting a phenotype of partial epithelial to mesenchymal transformation (EMT), which has been previously described in aggressive PDAC [35, 36].

**Figure 3.**
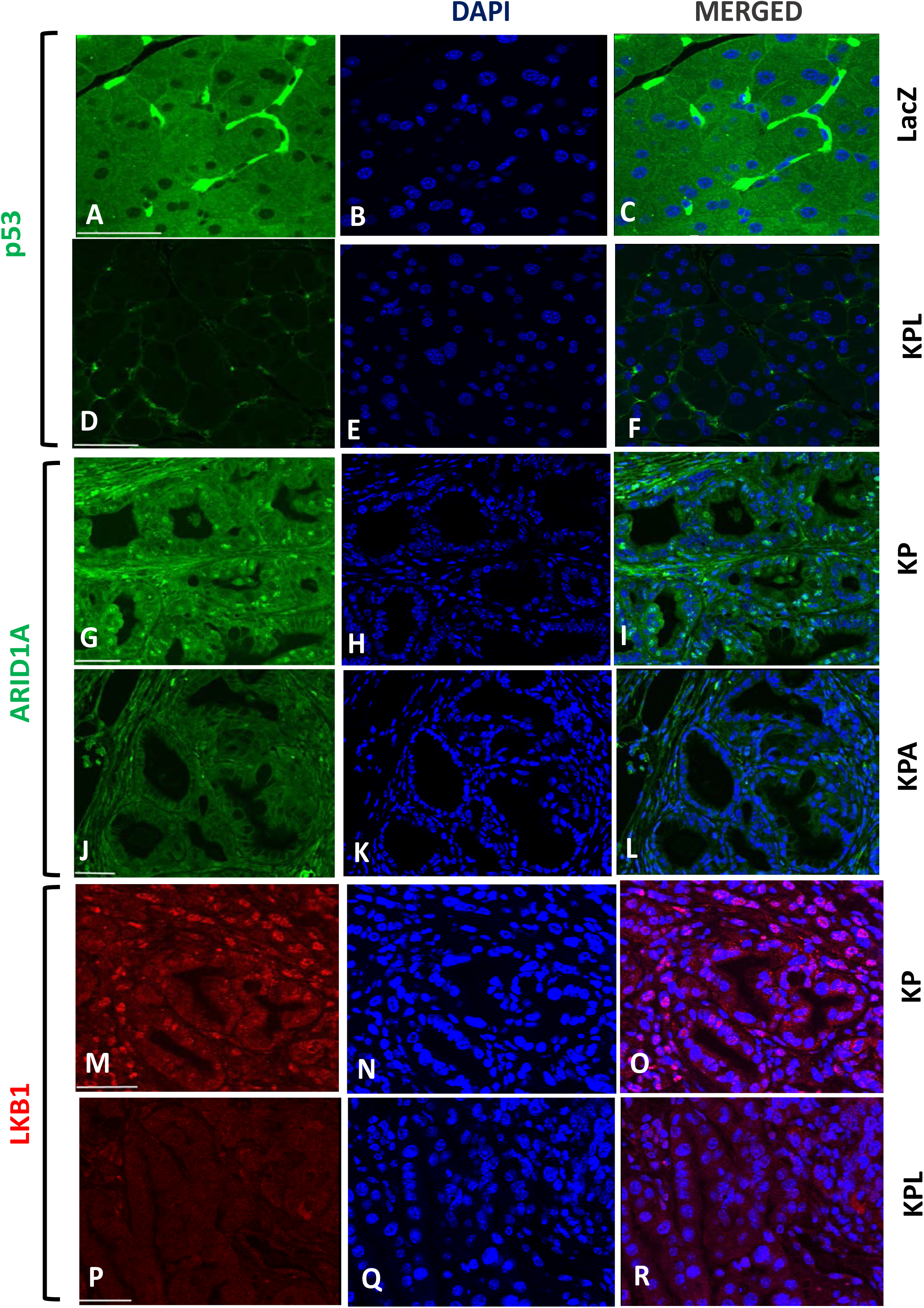
Downregulation of CRISPR/*Cas9* targeted protein in PDAC samples. **A-F.** Immunofluorescence was performed for p53 protein expression in control pancreas (*lacZ* injected) and “KPL” mouse tumors, showing absence of p53 in latter. **G-L.** Immunofluorescence for Arid1a shows loss of nuclear Arid1a expression in “KPA” tumors, with retained expression in “KP” tumors. **M-R.** Immunofluorescence for Lkb1 shows loss of nuclear Lkb1 expression in “KPL” tumors, with retained expression in “KP” tumors In this photomicrograph panel, (A) and (D) are p53, (G) and (J) are Arid1a, and (M) and (P) are Lkb1 stains, respectively. (B), (E), (H), (K), (N) and (Q) are DAPI stains. The scale bar = 50µ.

The T7 endonuclease assay serves as a preliminary measure of the efficacy of genome targeting based on the ability of the enzyme to cleave DNA strands that are not perfectly matched due to non-homologous end joining. Representative examples of the T7 endonuclease assay for *Kras*, *Trp53* and *Lkb1* from “KPL” and “KPA” tumors are shown in **Figure 4A**, which demonstrates targeting of sgRNA to the corresponding locus. In contrast to the cancer samples, the control pancreas injected with LacZ sgRNA showed no evidence of genome editing in the T7 endonuclease assay.

**Figure 4.**
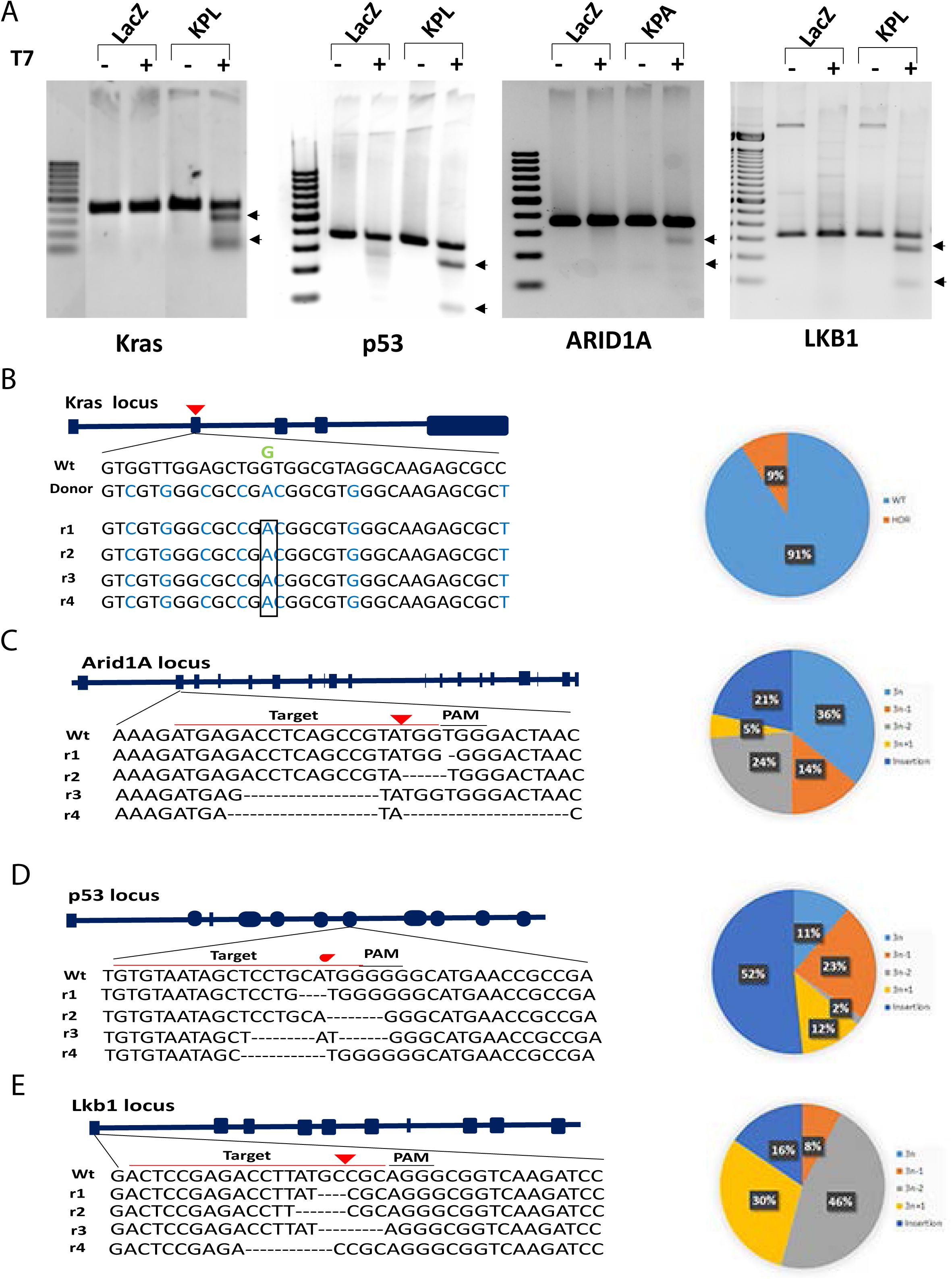
Sequencing of CRISPR/*Cas9* induced tumors confirms homology directed recombination (HDR) and gene editing at targeted loci. **A.**T7 endonuclease assay in genomic DNA isolated from “KPL” and “KPA” tumors following injection of AAV-sgRNAs constructs demonstrates evidence of gene targeting at the corresponding loci. Genomic DNA from AAV-LacZ injected pancreas was used as control. **B.**Schematic of sgRNA position and HDR donor template design for targeting the mouse endogenous *Kras* locus to incorporate a *Kras*^G12D^ oncogenic mutation. Next generation sequencing (NGS) reads confirms a G-A point mutation (*box*) leading to *Kras*^G12D^ while blue letters represent exogenous bases from the introduced HDR template that result in synonymous changes. The pie chart on the right represents the percentage of HDR at the *Kras* locus from the total number of NGS reads across all six of the sequenced mouse tumors. **C-E**. Schematic representing the sgRNA design for targeting *Arid1a*, *Trp53* and *Lkb1*, respectively, and representative NGS reads (r_n_) confirming the presence of “indels” that disrupt the target of interest (left). Notably, any given tumor demonstrated multiple indels at the targeted locus, consistent with a oligoclonal nature of the resulting neoplasm. The distribution of indel classes at each target site of interest is shown in the pie charts on the right.

In order to confirm the CRISPR/Cas9 mediated genome editing at the nucleotide level, we then performed exome sequencing of tumor derived genomic DNA from “KPL” and “KPA” virus injected pancreata, using reference DNA from a panel of C57/BL mice. In **Figure 4B**, we demonstrate NGS reads for *KRAS*, showing replacement of the endogenous locus with the donor template carrying the codon 12 non-synonymous mutation by HDR. Additional “marker” nucleotides (in blue color on sequence read) derived from the exogenous *KRAS* template (and accounting for synonymous base variations) are seen in the neoplastic DNA, confirming that the *Kras*^*G*^*12*^*D*^ mutation observed was not spontaneous but rather derived from the exogenous template by HDR. Since “bulk” tumor DNA was sequenced, we observed a mutant fraction of 9% for the *Kras*^*G*^*12*^*D*^ allele suggesting ∼20 % neoplastic cellularity. Exome sequencing data also confirmed disruption of three loci (*Arid1a*, *Trp53* and *Lkb1*), with multiple insertion deletion (“indel”) events observed for each gene (**Figures 4C, D and E**). This is consistent with independent CRISPR editing events occurring during the initial injection, and the observed PDAC “mass” at necropsy being of oligoclonal origin. The pie charts adjacent to each gene designate an overview of the classes of indel events for each targeted locus (detailed information for individual mouse tumors sequenced is illustrated in **Supplementary Figure 7**). The majority of indels were located 1-5bp upstream or downstream of the PAM sequence. The maximum insertion size observed was 15bp and the maximum deletion size observed was 23bp. Overall, between 8-30% of DNA reads (based on underlying neoplastic cellularity and gene of interest) demonstrated unequivocal indels on sequencing.

Finally, we explored whether the resulting tumors had developed secondary mutations of the 12 most commonly altered genes in PDAC, using data collated from the TCGA and ICGC datasets [3, 4]. Besides the CRISPR targeted genes in the corresponding mouse model (**Supplementary Figure 8**), we did not observe any secondary mutations in genes such as *Smad4*, *Gnas*, *Rnf43*, *Kdm6a* and *Brca2*. Importantly, the notable exceptions were *Kmt2c* (*Mll3*) and *Kmt2d* (*Mll4*), which encode for chromatin regulatory proteins [37], and demonstrated frameshift mutations or “indels” in subsets of tumors. It is unclear whether these events represented a requirement for tumor progression in these models, given that in the TCGA and ICGC datasets genes encoding for chromatin regulatory proteins are one of the most commonly mutated family of genes [3, 4].

## DISCUSSION

GEMMs represent the most rigorous preclinical platform for hypothesis testing the functional contribution of recurrent genetic alterations towards multistep pancreatic carcinogenesis, but conventional approaches demonstrate several logistical and temporal challenges, particularly, when complex allelotypes are desired. In this study, we have successfully demonstrated CRISPR-induced somatic gene editing using AAV-mediated direct intra-pancreatic injection of diverse sgRNA combinations for establishing autochthonous PDAC with high efficiency, and for some of the genotypes, relatively short latency to cancer incidence. Specifically, while five distinct sgRNA combinations were used as AAV cargo and all developed mPanINs, three genotypes (“KP”, “KPA”, and “KPL”) resulted in PDAC in the arbitrarily assigned time frame for observation (∼3 months), underscoring the importance of the genetic background on the latency of cancer formation. While it is certainly possible that the remaining two genotypes (“K” and “KA”) might eventually develop PDAC on longer follow up, our goal in this initial feasibility study was to keep the time frame for observation relatively short and perform necropsy on any surviving mice at ∼3 months. Importantly, all of the resulting PDAC lesions developed on a backdrop of credentialed precursor lesions (mPanINs and IPMNs), and progressed to metastatic disease in a subset of mice, confirming that the autochthonous models we have developed recapitulates the multistep progression seen in conventional *Cre*-based models. Loss of CRISPR-disrupted protein expression was confirmed in both pre-invasive and invasive neoplastic lesions, and NGS on the resulting tumors validated gene targeting at the sequence level.

The catalogue of genetic alterations targeted using CRISPR sgRNAs in the pancreas were based on the frequency of alterations reported in human PDAC in public databases [3, 4]. For example, *KRAS*, which is activated by oncogenic mutations in >90% of PDAC, and *TP53*, which is inactivated by loss of function mutations of homozygous deletions in 75% of PDAC, are obvious choices. In fact, mutations of *KRAS* are required not only for the cell autonomous effects in cancer progression, but also for the paracrine effects on the tumor microenvironment that create a tumor permissive immune milieu [38, 39]. Somatic mutations of the SWI/SNF complex *ARID1A* are seen in approximately 8-10% of PDAC, and this DNA binding helicase has been implicated in maintenance of acinar cell identity, and attenuating the progression to a precursor ductal phenotype (acinar ductal metaplasia) on a mutant *Kras* background [40, 41]. In our series, confirmed loss of *Arid1a* and expression of mutant *Kras* in “KA” mice was sufficient to induce mPanINs, but not PDAC within the ∼3 month time window. In contrast, addition of *Trp53* deletion (“KPA”) resulted in PDAC with relatively short latencies, with a greater propensity towards differentiated adenocarcinomas than the poorly differentiated carcinomas observed in “KP” mice alone. Of note, this correlation between *ARID1A* mutations and histological grade of differentiation has also been reported in other solid cancers, such as endometrial, hepatocellular, and pancreatic carcinomas [42-44], suggesting that loss of this epigenetic regulator might promote the acquisition of a ductal (i.e. “differentiated”) phenotype in PDAC. Finally, we also targeted Lkb1, which encodes for a serine threonine kinase mutated in several solid cancers. In the pancreas, *LKB1* mutations or loss of Lkb1 expression has been described in cystic precursors (IPMNs) [45, 46], and prior conventional models of conditional *Lkb1* loss have demonstrated PDAC arising on the background of cystic neoplasms [47, 48]. In our series, we did not examine “KL” as a dual targeting construct, but “KPL” injected mice developed cystic lesions resembling IPMNs in a subset of cases. Similar to “KPA” mice, “KPL” mice also developed predominantly adenocarcinomas upon progression.

A few important methodological caveats emerged during the course of this study. First, we recognized the AAV8 serotype to be significantly more efficient at viral transduction than AAV6, which is consistent with prior reports of using AAV8 as a delivery vector to the pancreas [49]. Second, the original CRISPR construct used in this study had a *Cre* expressing element [19], which resulted in spurious *Cas9* activation and extra-pancreatic mesenchymal tumors in the abdomen due to leakage of injected virus (*data not shown*), therefore, we deleted the *Cre* element, thus restricting recombinase expression to only the p48-expressing acinar parenchyma in the *p48-Cre*;LSL-*Cas9* mouse. Third, while there have been prior studies using CRISPR/*Cas9* for somatic gene editing and PDAC induction in mice, there is one key distinction from our approach. For example, Chiou et al [26] used an intraductal lentiviral injection methodology for delivering *Cre* recombinase and sgRNA against *Lkb1* into the main pancreatic duct of mice expressing conditional (“LSL”) allele of *Cas9* and mutant *Kras*^G12D^. Analogously, Maresch et al [27] used direct electroporation of plasmid based CRISPR/cas9 vectors against as many as 13 targeted loci in *Ptf1*-*Cre*; LSL-*Kras*^G12D^ mice, resulting in invasive tumors with multiple disrupted alleles. Both of these methods, however, required a conditional mutant LSL-*Kras*^G12D^ allele in the host animal (either constitutively expressed [27], or activated via virally delivered *Cre* [26]), on which background, somatic gene editing was enabled. In our study, however, we have dispensed with the requirement for a *Kras*^G12D^ allele in the host animal, by incorporation of a mutant template sequence in the AAV vector that is incorporated at the endogenous *Kras* locus through HDR. This has allowed us to utilize a single uniform strain of “workhorse” mice – *Ptf1-Cre*; LSL-*Cas9* – for injecting various sgRNAs, thereby, greatly enhancing the efficiency of PDAC generation when using different combinations of targeting vectors. Further, by expressing mutant *Kras* only in the subset of cells where the cooperating tumor suppressor allele is also disrupted, we ensure a *bona fide* “segmental” nature of disease in the pancreas, recapitulating the cognate human pathophysiology. A potential corollary advantage of this method is the possibility that genes of interest can be independently disrupted, including prior to the activation of mutant *Kras*, by varying the timing of AAV/CRISPR delivery (for example, with sequential injections).

Of interest, NGS studies on the CRISPR-induced PDAC showed that editing events were typically localized within a few bases of the PAM sequence, with individual “bulk” tumors harboring multiple classes of “indels” or frameshift mutations. This is consistent with the CRISPR-induced PDAC being an oligoclonal mix of more than one initiating clonal event, as previously reported [26]. While we did not pursue this particular line of investigation, one can envision that barcoding individual AAV constructs could allow for rigorous interrogation of selectivity in tumor formation and metastatic ability of distinct genomic alterations even at one given locus. The NGS data also revealed a somewhat robust rate of HDR for *Kras* in the sequenced “bulk” tumors (9% on average across six tumors), which is substantially greater than the 1.8% *Kras* HDR frequency reported in generating lung cancer models using AAV-mediated delivery of CRISPR [19]. The precise reason for this in unclear, but might represent a higher propensity for HDR in the pancreatic acinar component compared to the pulmonary epithelium, or a greater ability of the AAV8 serotype used in this study to enable HDR in the mouse genome, independent of exonuclease activity, as was recently shown for certain AAV strains [50]. Finally, our sequencing data showed that while several recurrently mutated PDAC-associated genes, including *Smad4*, *Gnas*, *Rnf43*, *Kdm6a* and *Brca2* did not undergo secondary mutations during PDAC development, two genes in particular - *Kmt2c* (*Mll3*) and *Kmt2d* (*Mll4*), which encode for chromatin regulatory proteins [37] – did demonstrate recurrent alterations. The encoded proteins, Mll3 and Mll4, are histone methyltransferase enzymes that are part of a so-called COMPASS-like complex responsible for addition of methyl residues to lysine residues on histones, which function as an “activation mark”. Loss of function mutations of *MLL3* and other COMPASS-like complex proteins is reported in approximately 10% of PDAC, and loss of expression has been associated with better prognosis [37]. We are continuing to explore the relevance of these recurrent non-targeted mutations, including their requirement for tumor formation in the context of other targeted loci using the AAV-CRISPR system.

In summary, we have generated a “workhorse” *Ptf1-Cre*; LSL-*Cas9* mouse that enables the relatively facile interrogation of the functional role of mutated genes in the PDAC landscape, including diverse combinations of targeted alleles using AAV8 as a delivery vector. Our platform precludes the need for a germline mutant *Kras* allele in the host mouse, and the resulting pancreata demonstrate the full compendium of precursor and invasive lesions observed in human PDAC. This “workhorse” platform will allow investigators to conduct *in vivo* functional genomics studies and generate reagents, such as cell lines with defined genetic alterations, with considerable efficiency.

## SUPPLEMENTARY FIGURES

**Supplementary Figure 1: Generation of Adeno-associated virus (AAV)-sgRNA constructs and evaluation of AAV vectors for intra-pancreatic injection.**

**A.** Schematic of various AAV constructs that were used in the study. GFP, green fluorescence protein; K, *Kras*; P, *Trp53*; A, *Arid1a*; L, *Lkb1*. KPLδCre denotes deletion of *Cre* from the parental KPL construct. **B-C.** Transduction of AAV6-Cre-GFP (**B**) and AAV8-Cre-GFP (**C**) by direct injection into the pancreas of adult B6 mouse. Diffuse GFP expression is shown in the injection site (pancreatic tail) at one month following injection with AAV8 subtype (**C**).

**Supplementary Figure 2: Representative examples of cystic papillary neoplasms consistent with intraductal papillary mucinous neoplasms (IPMNs) arising in the pancreata of “KPL” mice.**

The epithelial lining resembles high-grade pancreato-biliary subtype of IPMNs observed in humans.

**Supplementary Fig 3. Expression of Mucin1 in pancreatic precursor lesions of “KPL” mice.**

Low power (**A**-**C**) and high power (**D**-**F**) photomicrographs showing Mucin1 (green, **A**, **D**) expressed on the luminal surface of papillary epithelial cells in the IPMN-like precursor lesions. E-cadherin expression is shown in red **(B, E)** while DAPI in blue shown in the merged picture (**C**, **F**). Scale bar= 50µ.

**Supplementary Fig 4: Detailed histopathological assessment of subtypes of CRISPR-induced PDAC in AAV injected mice.**

**A**. On low power examination, the segmental nature of the invasive cancer is confirmed in CRISPR induced PDAC, with uninvolved pancreatic parenchyma to the left of the panel, and invasive neoplassia to the right. This contrasts with the usual diffuse nature of disease involvement in conventional Cre-driven models. **B.**The underlying genotype has an impact on the histology of the resulting PDAC in “KP”, “KPL’ and “KPA” mice. Thus, the majority of cancers in “KP” mice are poorly differentiated (sarcomatoid) carcinomas, while a smaller proportion are adenocarcinomas. In contrast, “KPA” and “KPL” both developed a higher frequency of conventional adenocarcinomas. Cases with no PDAC harbored high-grade PanINs. Of note, “KA” and “K” mice are not illustrated, as these did not develop cancers in any of the examined cases, although mPanINs and ADMs were observed.

**Supplementary Figure 5: Representative examples of metastatic PDAC in liver (A) and lung (B) of CRISPR injected mice.**

**Supplementary Figure 6: Immunofluorescence studies demonstrating loss of Arid1a protein in non-tumor bearing “KA” mice.**

**Upper panel.** Control mice with retained nuclear Arid1a expression (green) in native ductal epithelium (nuclear Sox9, red). **Middle panel.** Representative mPanIN lesion in a “KA” mouse pancreas, demonstrating presence of nuclear Arid1a in Sox9-expressing preneoplastic epithelium. **Lower panel.** Representative ADM lesion in a “KA” mouse demonstrating loss of nuclear Arid1a (green) and nuclear Sox9 (red); rare interspersed Arid1a retained cells are seen in the ADM lesion. (Scale bar: Upper panel = 50µ; Middle and Lower panel = 20µ).

**Supplementary Fig 7. Detailed tabulation of classes of insertion-deletion mutations (“indel”) and homology directed repair (HDR) in individual mouse tumor.**

**A-C.** Distribution and percentage of indel classes are shown for the *Arid1a* **(A),** *Trp53* **(B)** and *Lkb1* locus **(C)** from the six mouse tumors sequenced. **D.** The percentage of *Kras*^G12D^ HDR from the NGS reads in individual murine tumors.

**Supplementary Figure 8. A distribution of CRISPR-targeted and secondary alterations amongst the top 12 recurrently mutated genes in human PDAC**.

Overall, in the six murine tumors with NGS data, no secondary mutations are observed in most of the recurrently mutated genes in human PDAC, with the exception of *Kmt2c* and *Kmt2d*, which demonstrate frameshift mutations or indels in multiple tumors.

